# Network Pharmacognosy of *Galphimia glauca*: Mapping the Molecular Landscape of a Traditional Mexican Medicinal Plant

**DOI:** 10.1101/2025.01.02.631171

**Authors:** María José Cambero Acosta, Guillermo de Anda Jáuregui

## Abstract

This study explores the pharmacological landscape of *Galphimia glauca*, a traditional Mexican medicinal plant known for its sedative and anti-inflammatory effects. Using network pharmacognosy, we analyzed the interactions of *G. glauca*’s bioactive compounds, Galphimines A-I, with human protein targets. SwissTargetPrediction identified 214 unique protein targets across the galphimines, revealing a core-periphery structure in a bipartite network where 41 targets are shared among all compounds. Further interaction analysis using STRING-DB generated a dense protein-protein interaction network comprising 1,386 connections. Centrality analysis highlighted proteins such as SRC, MTOR, and MAPK3 as key nodes involved in cell growth, proliferation, and immune regulation pathways, suggesting these as pivotal mediators of *G. glauca*’s pharmacological effects. Community detection with the Walktrap algorithm further segmented the network into functionally relevant modules linked to cell survival, immune response, and inflammation, reflecting the therapeutic effects historically attributed to *G. glauca*. Our findings underscore the plant’s multi-target therapeutic potential and highlight the value of network-based approaches in understanding traditional medicine. This work lays the groundwork for further studies aimed at refining therapeutic strategies based on *G. glauca*’s bioactive compounds and suggests network pharmacognosy as a promising tool for assessing other traditional medicinal plants.

## Introduction

*Galphimia glauca* is a widely used plant in traditional Mexican medicine, recognized for its therapeutic sedative and tranquilizing effects (1). This shrub, commonly referred to as red arnica, belongs to the Malpighiaceae family and grows between one and three meters in height. Characterized by clusters of yellow flowers, *G. glauca* has ovate or elongated leaves that are green on top and bluish underneath. It is a native species in Mexico, thriving in mountainous regions as well as in flatlands across central and northeastern states such as Aguascalientes, Guanajuato, and Jalisco, where it grows wild(2).

The bioactive compounds of *G. glauca*, known as Galphimines A-I, are responsible for its various medicinal properties, including antioxidant, antiproliferative, proapoptotic, antiangiogenic, and anti-inflammatory activities (3). The antioxidant properties help inhibit oxidative processes, reducing free radical production. Its antiproliferative effects suppress the growth of cells, particularly cancer cells, through pathways like JAK and STAT. Proapoptotic effects induce programmed cell death, helping eliminate damaged cells. The antiangiogenic activity inhibits the formation of new blood vessels, primarily through interactions with tyrosine kinases, while anti-inflammatory effects modulate immune responses, addressing chronic inflammation linked to diseases such as arthritis and neurodegenerative disorders.

In recent years, understanding the complex interactions of medicinal compounds has become increasingly important in developing safe and effective treatments. *Network pharmacognosy* applies network science to the pharmacology of natural products, examining how multiple compounds in a medicinal plant, like *G. glauca*, interact with various molecular targets and pathways. This approach helps visualize the interconnected effects of *G. glauca*’s compounds on human health, providing insights into its medicinal applications and guiding the development of more precise therapeutic agents without adverse side effects.

## Results and discussion

### Identification of Pharmacological Targets

The bioactive compounds of *Galphimia glauca*, known as Galphimines (A-I), were analyzed using the SwissTargetPrediction tool to identify their potential pharmacological targets. The results identified a total of 180 distinct protein targets across all galphimines, with varying probabilities of interaction. A comprehensive set of selected interactions for all targets can be found in Supplementary material 1.

### Target Space of Galphimines

The target space of *Galphimia glauca* was analyzed, revealing a single, connected component encompassing 214 protein targets identified for the galphimines. A significant level of target overlap was observed, with 41 targets shared across all galphimines. This indicates potential common pathways or biological mechanisms influenced by the different compounds within *G. glauca*. A visualization of this target space may be found in Figure 1.

**Figure 1:**
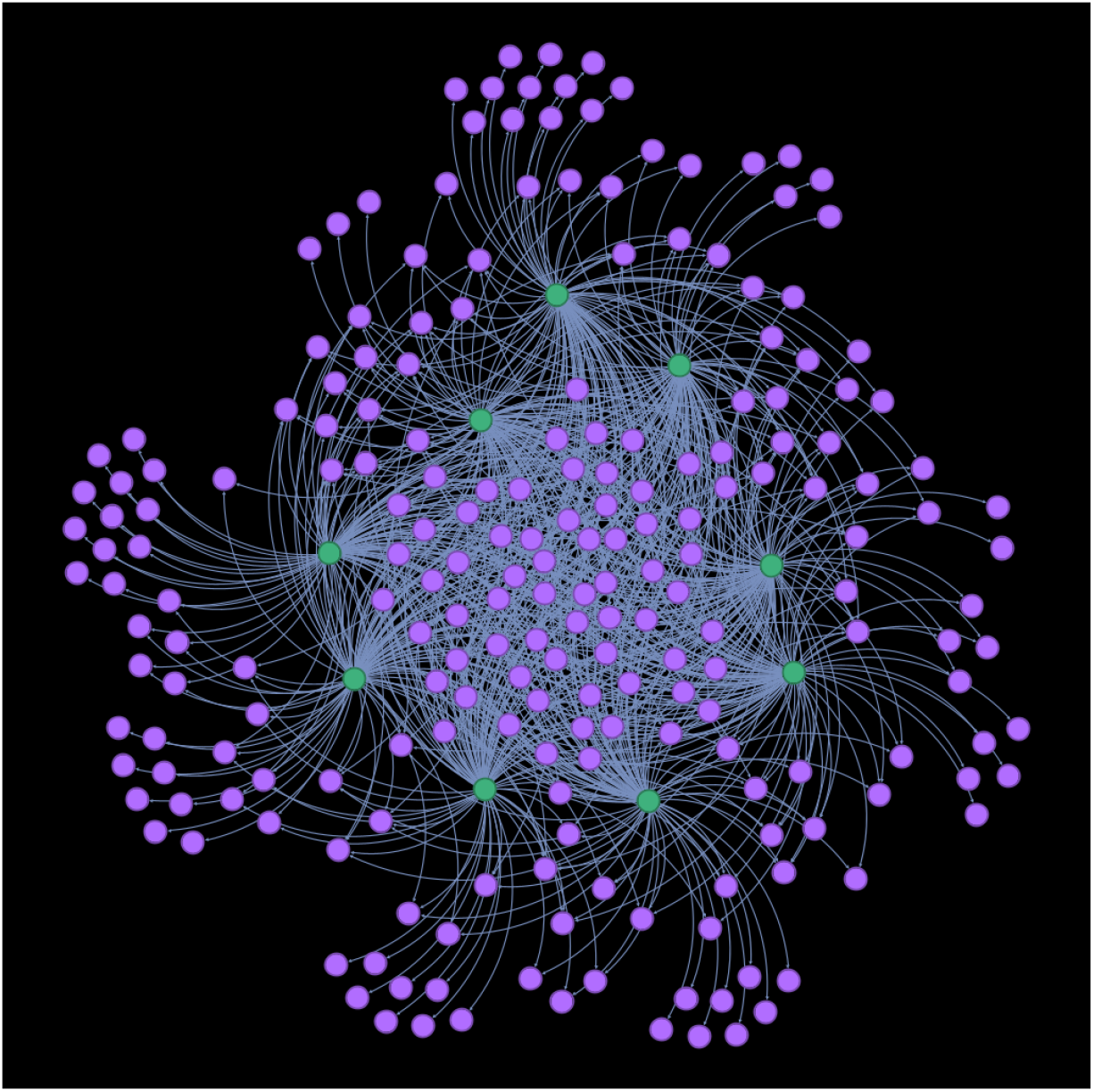
network visualization of the target space of *G. glauca*, represented as a bipartite network. Galphimines are shown in green, and molecular targets in purple. A core-periphery structure may be observed in the network.

We represented this target space as a bipartite network anchored by a **core structure** of 41 shared targets surrounded by a **peripheral layer** of 63 unique targets, each associated with a single galphimine. The degree distribution of these targets, shown in Figure 2, reflects their connectivity within the network. Full descriptors and data for all targets are available in Supplementary material 2.

**Figure 2:**
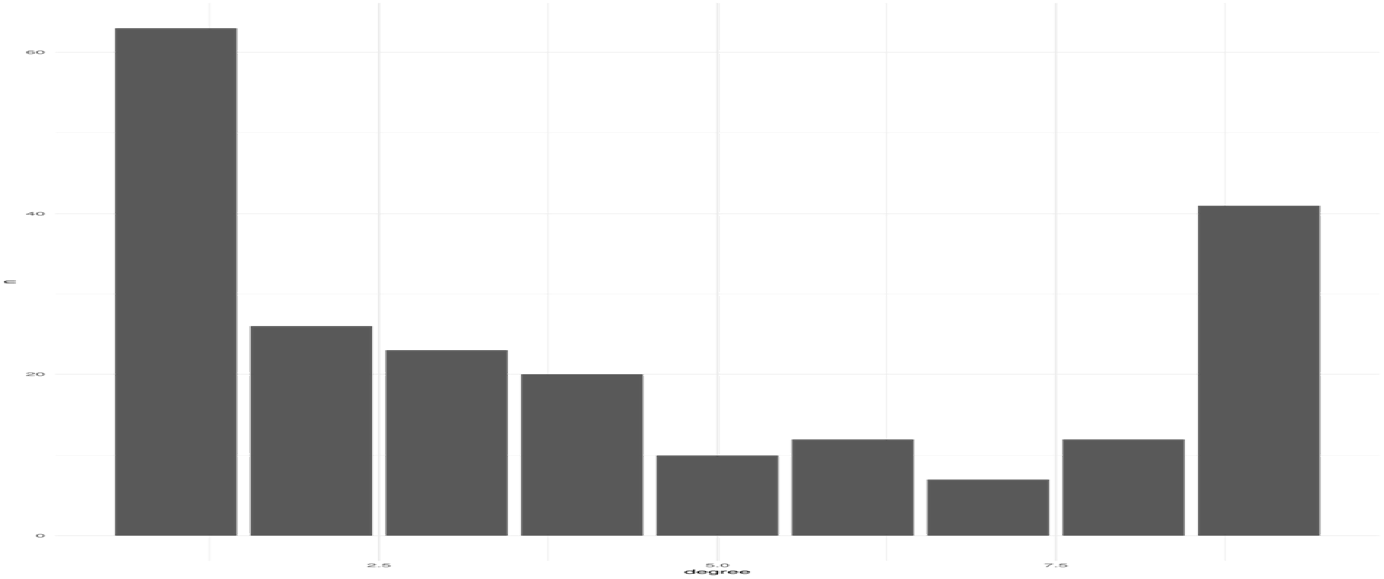
Degree distribution of target proteins in the *G. glauca* target space. Notice that the core structure (nodes connected to all galphimines) and the periphery structure (nodes connected to a single galphimine) are more abundant in the target space.

Biologically, several targets within the core structure point to possible pharmacological activities of *G. glauca*. Noteworthy proteins such as Protein Kinase C Alpha (PRKCA), Glucocorticoid Receptor (NR3C1), and P-Glycoprotein 1 (ABCB1) were identified across multiple compounds, suggesting that these shared targets could contribute to the therapeutic effects reported for *G. glauca*.

### Protein Interaction Network of *G. glauca* Targets

We mapped the pharmacological targets of *G. glauca* using STRING-DB, revealing a network of 1,386 interactions. This network provides insights into how these targets interact within their biological contexts, positioning each protein’s role in a complex web of signaling pathways. A visualization of this network is shown in figure 3. The full network is provided in graphml format as Supplementary material 3.

**Figure 3:**
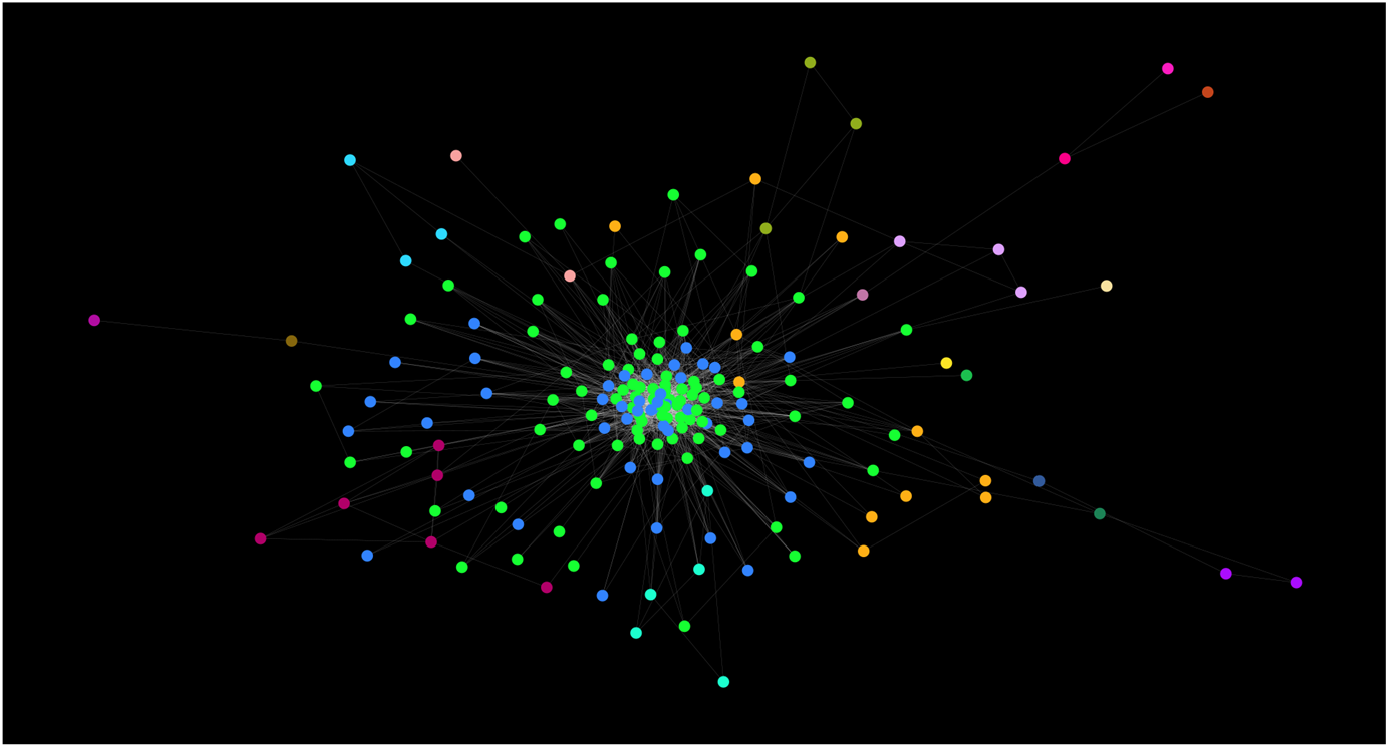
Network visualization of protein-protein interactions between targets of *G. glauca*. Nodes are colored according to the community they belong as determined by the Walktrap algorithm.

In figure 4, we show the value distribution of centrality measures in this network. By the distributions, we may observe a *heavy-tailed* behavior for both degree centrality and betweenness centrality.

**Figure 4:**
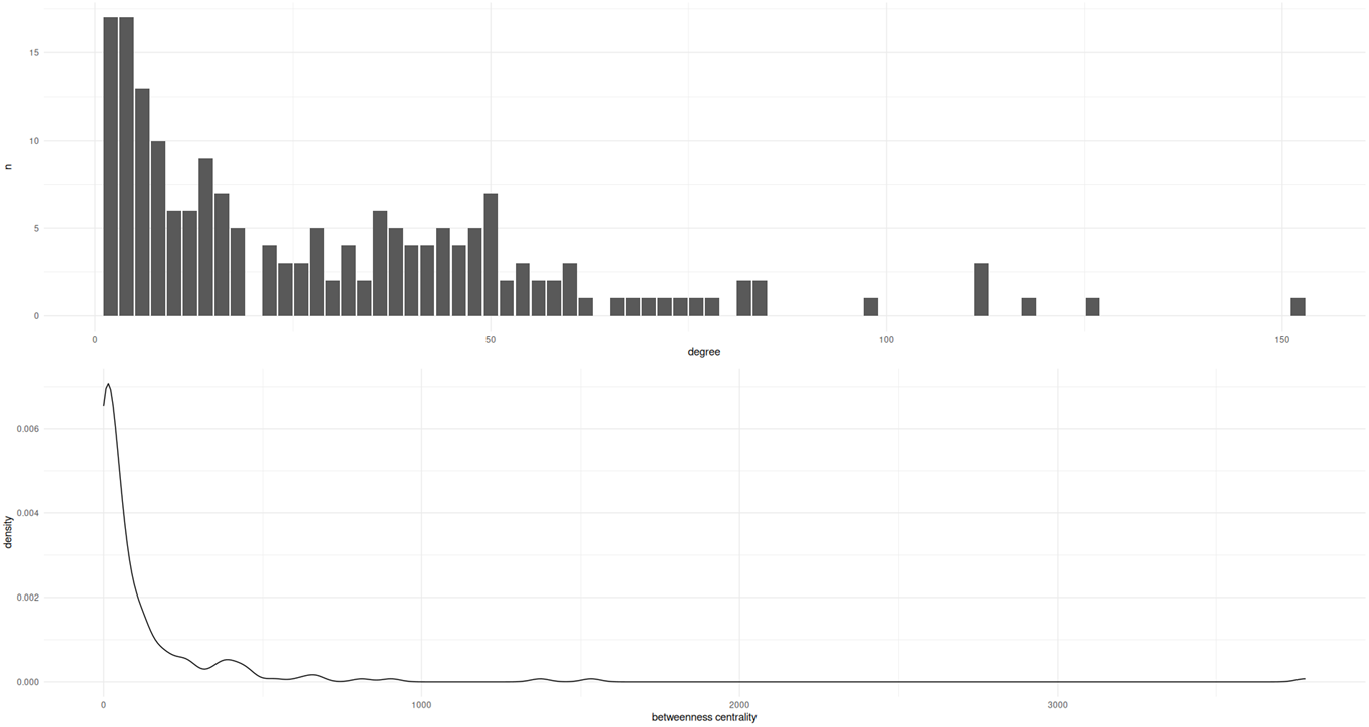
Distribution of centrality measures on the interaction network of *G. glauca* targets. Top panel shows histogram of degree centrality, while bottom panel shows density plot for betweenness centrality. In both cases, heavy-tailed distributions are shown, indicating the emergence of central elements in the network.

As such, certain proteins emerged as key players, their centrality measures underscoring their influence. Degree centrality identifies SRC, MTOR, MAPK3, and MAPK1 as the most connected proteins, each prominently involved in pathways crucial for cell growth, proliferation, and survival. Their connectivity suggests these proteins are central mediators of *G. glauca*’s pharmacological activity. Betweenness centrality, which highlights nodes that bridge different network regions, ranks SRC as the top communication hub. Other proteins with high betweenness centrality include JUN, PIK3CA, HSP90AB1, ADRB2, and MMP1, each influencing transcription regulation, signal transduction, or immune response. The whole set of network descriptors may be found in Supplementary material 4.

By looking at these individual proteins, we may gain further insight on the potential mechanisms through which *G. glauca* exhibits its known medicinal properties, as well as providing insights on possible future therapeutic applications. For instance:

- **SRC** supports immune processes, including proliferation, phagocytosis, and immune cell development. It advances cell cycle progression through interactions with receptors and ligands involved in macrophage-mediated inflammation, playing a recognized role in neoplastic disease treatments (4,5).
- **MTOR** is a critical metabolic regulator, modulating growth factors, hormones, and cellular stress responses while inhibiting autophagy. MTOR-targeted therapies show promise not only in cancer but also for neurodegenerative and metabolic disorders (6,7). Inhibiting the PI3K/Akt/MTOR pathway is an emerging therapeutic strategy in oncology (8).
- **PIK3CA** plays a key role in cell morphology and survival through growth factor responses. Notably, mutations in PIK3CA are linked to therapeutic resistance in about 30% of early-stage HER2+ tumors (9,10).
- **MAPK3** is involved in processes such as cellular migration, transcription regulation, and inflammation, particularly through TNF signaling (11).
- **JUN** is a transcription factor binding to the AP1 consensus sequence, facilitating T-cell-induced cell death and steroidogenic gene expression (12).
- **HSP90AB1** functions as a molecular chaperone, essential for the regulation, maturation, and structural maintenance of target proteins. Its ATPase-dependent cycle aids in epigenetic regulation and gene expression control (13).
- **ADRB2** mediates adenylate cyclase activation through epinephrine binding, with heightened affinity for norepinephrine. This receptor is also studied in prostate cancer therapy (14,15).
- **MMP1** exhibits catalytic activity on collagen types I, II, and III, contributing to inflammatory responses and playing a role in neuroinflammation, especially in HIV contexts (16).

### Community Structure

Applying the Walktrap algorithm to the network, we identified clusters of highly connected proteins, each linked to specific biological functions. Three main communities emerged, each aligning with distinct pharmacological activities of *G. glauca*:

- **Community 1** includes proteins like MTOR, MAPK3, and JUN, which play roles in apoptosis and cell survival. This aligns with *G. glauca*’s reported antiproliferative effects.
- **Community 2** encompasses SRC, PRKCA, and HSP90AB1, central to cell growth and immune response pathways, highlighting *G. glauca*’s role in cellular regulation and immune adaptation.
- **Community 3** is associated with inflammation and immune modulation, containing targets like ADRB2 and MMP1, and supporting the traditional anti-inflammatory uses of *G. glauca*.

As such, we may observe that the biological effects of *G. glauca* may not be only due to direct action on single molecules, but by a concerted modulation of several proteins acting in a coordinated manner.

## Conclusions

This study mapped the pharmacological target landscape of *Galphimia glauca* using network pharmacognosy, identifying 214 protein targets and analyzing their interactions. Proteins like SRC, MTOR, and MAPK3 emerged as central nodes in pathways governing cell growth, proliferation, and immune regulation. related to cell growth, proliferation, and immune regulation. Network analysis showed that *G*.*glauca’*s bioactive compounds do not act in isolation but instead target clusters of interconnected proteins. These clusters—centered on cell survival, immune response, and inflammation—align with *G. glauca*’s known medicinal effects, indicating that its therapeutic properties may result from coordinated activity across multiple pathways.

The results underscore the potential of *G. glauca* as a multi-target therapeutic agent and demonstrate the value of network approaches in understanding traditional medicines. By identifying how *G*.*glauca*’s compounds interact with key biological networks, this work provides a foundation for further investigation into its pharmacological effects and supports the development of targeted therapeutic strategies that leverage its traditional uses. Future studies may refine these insights, focusing on specific pathways to validate and extend *G. glauca*’s pharmacological effects. Furthermore, we believe that network pharmacognosy approaches may be of help to integrate other mexican traditional products for medical applications.

## Methods

### Selection of Bioactive Compounds

The pharmacologically relevant compounds from *Galphimia glauca*, known as Galphimines A-I, were extracted from the BIOFACQUIM database maintained by UNAM (17). BIOFACQUIM is a curated repository that aggregates high-confidence compound-target associations from nine public databases. This database applies a standardized curation protocol involving duplicate removal and molecule cleaning, ensuring data reliability and coverage across various biological contexts. The database may be accessed at: https://www.difacquim.com/d-databases/

The SMILES (Simplified Molecular Input Line Entry System) strings for each galphimine were obtained from BIOFACQUIM and verified against the NCBI database (https://www.ncbi.nlm.nih.gov/). These notations allowed for consistent data entry into prediction tools for further analysis.

### Identification of Pharmacological Targets

To identify potential protein targets, we used SwissTargetPrediction (18), (accessible at http://www.swisstargetprediction.ch/help_mol.php). This tool predicts probable protein targets based on ligand similarity in both 2D and 3D conformations. Predictions were limited to the Homo sapiens species to ensure relevance for potential therapeutic applications.

The SMILES strings for each galphimine were entered into SwissTargetPrediction using default parameters, and predicted targets were ranked by probability. Targets with the highest predicted affinity were retained for network analysis, resulting in a comprehensive list of 180 unique protein targets across the nine galphimines. The top 100 highest ranked targets for each galphimine were integrated into a single bipartite network representing the pharmacological target space for *G. glauca*.

### Molecular Interaction Search

Protein targets identified through SwissTargetPrediction were mapped for potential interactions using the STRING database (19) (accessible at https://string-db.org/). STRING aggregates protein-protein interaction data from a variety of sources, including computational predictions, knowledge transfer from model organisms, and experimental data. Default parameters were used, with Homo sapiens set as the reference organism.

### Network Analysis

To analyze the networks’ structures and highlight key nodes, we employed several centrality metrics:

Degree Centrality: Degree centrality was calculated as the number of direct interactions (edges) for each node (protein). Nodes with high degree centrality, such as SRC and MTOR, were flagged as potential hubs within the network.

Betweenness Centrality: Betweenness centrality was calculated using Freeman’s formula to determine each node’s importance in maintaining shortest-path connectivity across the network. This metric highlighted key bridging proteins, such as SRC and JUN.

Community Detection: We used the Walktrap algorithm to identify community structures within the network. Walktrap detects communities by simulating random walks on the network, under the principle that shorter random walks are more likely to remain within the same community. The algorithm iteratively merges nodes into communities by minimizing the mean distance between nodes within each community, creating a hierarchical structure of nested communities.

### Statistical and Visual Analysis

All network and statistical analyses were conducted in R (version 4.1.3). Network analysis was performed using the *igraph* package. Visualization of network topologies and centrality distributions were generated using *ggplot2*. Other visualizations were generated using the *gephi 0*.*10* software.

## Notes

### Competing Interest Statement

The authors have declared no competing interest.

